# BOLD differences normally attributed to inhibitory control predict symptoms, not task-directed inhibitory control in ADHD

**DOI:** 10.1101/699728

**Authors:** Andre Chevrier, Russell J. Schachar

## Abstract

**Background:** Altered brain activity that has been observed in attention deficit hyperactivity disorder (ADHD) while performing cognitive control tasks like the stop signal task (SST), has generally been interpreted as reflecting either weak (under-active) or compensatory (over-active) versions of the same functions as in healthy controls. If so, then regional activities that correlate with the efficiency of inhibitory control (*i.e.* stop signal reaction time, SSRT) in healthy subjects should also correlate with SSRT in ADHD. Here we test the alternate hypothesis that BOLD differences might instead reflect the redirection of neural processing resources normally used for task-directed inhibitory control, toward actively managing symptomatic behavior. If so, then activities that correlate with SSRT in TD should instead correlate with inattentive and hyperactive symptoms in ADHD.

**Methods:** We used fMRI in 14 typically developing (TD) and 14 ADHD adolescents performing the SST, and in a replication sample of 14 healthy adults. First we identified significant group BOLD differences during all phases of activity in the SST (*i.e.* warning, response, reactive inhibition, error detection and post-error slowing). Next, we correlated these phases of activity with SSRT in TD, and with SSRT, inattentive and hyperactive symptom scores in ADHD. We then identified whole brain significant correlations in regions of significant group difference in activity.

**Results:** Only three regions of significant group difference were correlated with SSRT in TD and replication groups (left and right inferior frontal gyri (IFG) during error detection, and hypothalamus during post-error slowing). Consistent with regions of altered activity managing symptomatic behavior instead of task-directed behavior, left IFG correlated with greater inattentive score, right IFG correlated with lower hyperactive score, and hypothalamus correlated with greater inattentive score and oppositely correlated with SSRT compared to TD.

**Conclusions:** Results are consistent with stimuli that elicit task-directed integration of neural processing in healthy subjects, instead directing integrated function towards managing symptomatic behavior in ADHD. The ability of the current approach to determine whether altered neural activities reflect comparable functions in ADHD and control groups has broad implications for the development and monitoring of therapeutic interventions.

## Background

Attention deficit hyperactivity disorder (ADHD) is associated with cognitive difficulties, particularly in the form of decreased inhibitory control (1,2), and with restless, inattentive and impulsive behavior compared to typically developing (TD) individuals. Inhibitory control, critically reliant on brain networks involving right inferior frontal gyrus (IFG) and caudate nucleus (3,4), can be measured using the stop signal task (SST) (5). The SST consists of a brief warning stimulus followed by a primary choice reaction time task, and an occasional (eg 33% of trials) stop task in which a stop signal is presented at some delay after the go stimulus. The stop signal delay adapts to performance, increasing after successful stop trials, and decreasing after unsuccessful stop trials. This adaptive delay ensures that only half of stop trials can be successfully inhibited on average. Stop trials that cannot be stopped generate performance errors in the form of responses that should not have been made. The SST can estimate the unobservable speed of stopping, called stop signal reaction time (SSRT) by subtracting the mean stop delay from the mean response time on trials with no stop signal (6).

Neuroimaging studies of ADHD using the SST have consistently found deficient inhibitory control to be associated with altered activity and connectivity (7–9). For the most part, altered activity and connectivity have been interpreted as evidence of either relatively weak (decreased activity/connectivity) or compensatory (increased activity/connectivity) versions of normal function. However, there has been little consideration given to the possibility that activity differences in ADHD might instead reflect altered integration of the same neural processing resources toward different goals rather than simply under- or over-activation. This distinction is important because etiological and therapeutic models implicated by under- or over-activation, which would aim to target and adjust specific functions, would differ from those implicated by differences in integration, which might instead aim to desensitize patients to the distracting effects of contextual cues on attentional control.

One indication that altered activities in ADHD do not simply reflect under- or over-activation is observations of opposite activity changes with respect to baseline compared to healthy control subjects. We found four such patterns of opposite activity in a recent study of ADHD and typically developing (TD) adolescents performing the SST. First, we observed opposite activity in task-related (deactivation instead of activation) and default mode networks (activation instead of deactivation) during response phases indicative of categorically altered preparation (9), Second, we noted opposite activity in ventral striatum (activation instead of deactivation) during post-error slowing, which correlated with heightened amygdala activity in ADHD instead of dorsal striatum as in TD (10). Opposite activation of ventral striatum and heightened activation of the amygdala during reward and prediction error processing have consistently been observed in ADHD (11–18). Heightened amygdala input to the ventral striatum prevents the kind of thresholding influences from ventral to dorsal striatum required for reinforcement learning (19) by enhancing limbic and motor processing while suppressing cognitive processing (20–22). Third, ADHD was characterized by opposite responses in non-dopaminergic nuclei (locus coeruleus, raphe and medial septal nuclei) during post-error slowing indicative of a categorically altered competition for control of dopamine (10). Neurotransmitter systems that compete for control of dopamine strongly influence the kind of attention given to stimuli (*e.g.* internally *vs.* externally directed (23–25)) and the learning generated by the outcome of a given trial (*e.g.* controlled reinforcement learning from task-related feedback *vs.* surprise and learning about environmental context (23–26)). Fourth, we saw opposite hypothalamus activity (activation instead of deactivation) and altered correlation of hypothalamus activity with reciprocally connected (27–29) neurotransmitter nuclei (substantia nigra, locus coeruleus, raphe and medial septal nuclei) during post-error slowing (10). The hypothalamus is a motivation-cognition interface for the control of integrated functions such as food-seeking and non-specific consummatory behaviors (30,31). The hypothalamus can orchestrate complex behaviors by mobilizing information processing in downstream targets and directing distributed processing resources towards unified goals, while suppressing processes associated with competing goals (32). Altered functioning of the hypothalamus could therefore strongly influence the integration of distributed neural processing and the goals to which they are directed.

Rather than reflecting relatively weak or compensatory versions of normal function, we propose that altered activity in ADHD might instead reflect altered integration of distributed neural processing resources toward non task-related goals. Processing resources that are used for effective task performance in healthy subjects might instead be directed toward supporting or suppressing symptomatic behaviors like wandering attention and impulsive behavior in ADHD. If so, then neural differences would be more analogous to optical rivalry (33) than to weak or compensatory function.

One form of optical rivalry is apparent when viewing bistable images, in which the contours of an image can be perceived as two distinct objects (eg. duck or rabbit) but both cannot be perceived simultaneously, because the dynamics of component neural processing resources (*e.g.* edge detection) can only represent one unified object at a time. Similarly, when subjects perform the SST, their component neural processing resources can only support one integrated focus of attention at a time. For example, activity in left inferior frontal gyrus, which has been consistently found to be altered in ADHD (34), is involved in suppressing interference (*e.g.* in the form of added noise or distracting stimuli) with efficient performance on a variety of tasks (35–38). Left inferior frontal gyrus invariably performs interference suppression, regardless of what is considered a noise and what is considered a signal, just as edge detection regions invariably perform edge detection regardless of the integrated object that is perceived (*e.g.* duck or rabbit). Although the stimuli used as interference in interference suppression tasks are under objective experimental control, what constitutes interference to our objective neural processing is in fact our subjective internal state, in the way that a meal might be represented in the brain as a signal, but leftovers as a noise.

We propose that in the kind of distracted and impulsive states associated with ADHD, the task itself might be processed as noise rather than signal. If so, then activities in regions that normally predict improved performance (*i.e.* SSRT) should instead predict symptoms in ADHD. For instance, neural activities in left and right inferior frontal gyri that, in typically developing individuals, perform inhibitory control and interference suppression processing, might, in individuals with ADHD, be directed toward suppressing impulsive behavior and supporting wandering attention. This distinction is crucial for the development of appropriate neurocognitive models that are increasingly being used to inform and monitor therapeutic interventions.

Here we attempt to determine whether altered activity in ADHD reflects either a) weak or compensatory versions of normal function, or b) the same neural processing resources managing wandering attention and hyperactive behavior instead of supporting efficient task performance. We test this hypothesis using the straight forward approach of examining the actual correlates these activities, which to our knowledge, has not been done before.

First we perform intersubject correlation analyses on all phases of SST activity (*i.e.* warning and response phases on all trials, response cancellation phases on successful stop trials, and error detection and post-error slowing phases on failed stop trials, reported in (9,10)) with SSRT in both groups, and with inattentive, hyperactive and total symptom scores in ADHD. Secondly, we inspect regions of significant group BOLD difference for whole brain corrected correlations with SSRT in TD. Given the concerns with replication in fMRI (39,40), we perform the same analyses in an independent replication sample of healthy adults, and only report TD correlations with SSRT that are present in both samples. Thirdly, we inspect regions of significant BOLD difference for whole brain corrected correlations with SSRT, inattentive, hyperactive and total symptom scores in ADHD. If deficient inhibitory control in ADHD is the result of relatively weak or compensatory versions of normal function then the same regions that predict SSRT in TD should also predict SSRT in ADHD. However, if altered activities are instead the result of deploying overlapping neural resources toward managing symptomatic behavior, then activities which correlate with SSRT in TD and replication groups should instead correlate with inattentive and hyperactive symptoms. Further, if ADHD activities that correlate with symptoms instead of SSRT are both not directed towards, and are actively directed against efficient task performance, then we would predict that these activities should be oppositely (*e.g.* negatively *vs.* positively) correlated with SSRT compared to TD.

## Methods

### 2.1 Subjects

This study is the third stage of analyses performed on data from TD and ADHD adolescents presented in (9,10). Fourteen adolescents diagnosed with ADHD (7 male, 12-17 years) and 14 TD adolescents (9 male, 12-17 years) were included in this study. Subjects gave informed, written consent and the study was approved by the Hospital for Sick Children institutional research ethics board. Written informed consent was obtained from the parents of all participants under the age of 16. ADHD subjects on stimulant medication (n = 6) stopped taking medication 24 hours prior to the scan to eliminate drug-induced BOLD changes (41).

ADHD subjects and their parents were interviewed separately and together using the parent interview for child symptoms (PICS-IV (42)). Intelligence was assessed using the Wechsler Intelligence Scale for Children (WISC-IV). ADHD subjects met diagnostic and statistical manual of mental disorders (DSM-5) criteria for ADHD (at least six out of nine inattentive symptoms, hyperactive-impulsive symptoms, or both according to at least two of three informants (parents, teacher and/or patient self-report)). ADHD subjects also showed moderate to severe impairment in both school and home settings (Global Assessment Scale (43) score < 60). Subjects were excluded if they had any comorbid psychiatric or neurological disorder other than oppositional defiant disorder (ODD) or learning disability within the previous 12 months (*e.g.*, obsessive compulsive disorder, Tourette syndrome, major depressive, anxiety or pervasive developmental disorder), an IQ score of below 80 on verbal and performance scales or any medical issues that would impact fMRI participation. Subjects with contraindications for MRI (metal braces or metal fragments in their body) were also excluded.

Nine ADHD subjects were diagnosed with ADHD combined subtype, five met criteria for inattentive subtype, and two also met DSM-5 criteria for ODD. Control subjects were assessed in a comparable manner and reported no psychiatric or medical disorders. All subjects were right-handed and had normal vision and hearing.

The replication sample used for comparing results of behavioral correlations in TD consisted of 14 healthy young adults on a placebo dose of methylphenidate (8 male, mean (±SD) age = 24.0 ±2.8 years). Subjects from the replication sample gave informed, written consent and the study was approved by the Hospital For Sick Children institutional research ethics board. These data also served as the replication sample in our previous paper examining error processing activities (10).

### 2.2 Behavioral task

The stop signal task (SST) (44) involves a primary choice reaction time task and a secondary stop task. Trials began with a fixation point in the centre of a black screen (500 ms), followed by the go-stimulus (1000 ms). Subjects were instructed to respond as quickly and accurately as possible with their left thumb when the letter “X” appeared or with their right thumb when the letter “O” appeared. In 33% of trials, a stop signal (background colour change from black to red) followed the go stimulus. Subjects were instructed to stop if they saw the stop signal, but not to wait for stop signals. The initial stop signal delay was 250 ms and increased/decreased by 50 ms after successful/unsuccessful stop trials, ensuring 50% stop errors on average. The task involved 224 trials, requiring a total scan time of 15 minutes.

Inter-trial interval (ITI) was jittered to maximise the number of independent equations in the deconvolution analysis, using trials of 2.5 or 3.5 seconds. Every fourteenth trial was followed by a 17.5 second rest (blank screen). Trial order was pseudo randomised so that the current trial did not predict the subsequent type of trial. Mean go response time (RT) was observable from the 67% of trials in which no stop signal appeared. Stop signal reaction time (SSRT) was estimated by subtracting the mean delay on stop signal trials from the mean RT on trials with no stop signal. Behavioral scores (within-group means and between-group differences) were analysed using two-tailed t-tests.

### 2.3 Scanning Parameters

Imaging was done with a GE LX 1.5T MRI scanner (GE Healthcare, Waukesha, WI). Anatomical data were acquired with a standard high-quality SPGR sequence (120 slices, 1.5 mm thick, FOV 24 cm, 256 × 256 matrix). Functional data were collected using a GRE-EPI sequence with an 8-channel head coil (TE = 40; TR = 2,000; Flip angle =90 degrees; 24 slices; 6 mm thick; FOV 24 cm; 100 kHz readout bandwidth; 64×64 in-plane resolution). Behavioral data were collected using a fiber-optic response system interfaced to a laptop running the SST.

### 2.4 Single subject analysis

Functional data were analysed using AFNI version 16.0.09 (45). Images were motion corrected and inspected to ensure motion did not exceed 3 mm or 3 degrees. We used a standard motion correction algorithm and censored noisy time points (>3.5 median absolute deviations). We used a general linear model of stimulus vectors convolved with the hemodynamic response function (HRF) using AFNI’s 3dDeconvolve program. Estimates of baseline (seventh order polynomial) were generated along with 6-point HRF’s for all event types (HRF delay = 2TR).

The following event types were used in the deconvolution analysis (as in (4)): fixate (F), time-locked to warning-stimuli at the beginning of every trial, left- (X) and right-hand (O) response events, time locked to motor responses, successful inhibition (SI), time-locked to the presentation of stop signals, and error detection (Detect) and post-error slowing (PES) events, both time-locked to responses on failed stop trials. Go trials were modelled using (F) and (X) or (O) stimuli. Proactive inhibtion activity during response phases was isolated using the contrast (X+O)/2 in order to suppress hand-specific response activity from relatively hand-independent proactive inhibition as in (4,9,47). Successful stop trials were modelled using (F) and (SI). Activity during reactive inhibition was identified using the contrast SI - (X+O)/2 as in (4,9). Failed stop trials followed by less than median response slowing were modelled with (F), (X) or (O), and (Detect). Failed stop trials followed by greater than median response slowing were modelled with (F), (X) or (O), (Detect) and (PES). Activation maps were estimated by taking the area under the HRF, warped into Talairach space (1mm^3^ resolution), and smoothed using a 6mm full width at half maximum Gaussian kernel.

### 2.5 Group level ANOVA analysis

Single subject activation maps were entered into a random effects ANOVA analysis for ADHD and TD groups. Statistics for group wise ANOVA output maps were distributed as a t* statistic with 13 degrees of freedom. Group difference maps were generated using a nested repeated measures 3-factor ANOVA (group membership, event types, and subjects) to identify significantly different activities between TD and ADHD adolescents. Statistics for group difference ANOVA output maps were distributed as a t* statistic with 26 degrees of freedom. Output from all ANOVA analyses (TD, ADHD, TD-ADHD) were corrected for multiple comparisons using AFNI’s 3dClustSim program (48) (spatial correlation estimated from AFNIs corrected 3dFWHMx -acf option). This analysis required significant voxels to be part of a cluster of at least 10.9 original voxels (920 mm^3^) with a minimum Z score of 1.96 for an overall α < 0.05. The low threshold and large cluster size used for correction is based on the expected effect sizes from our previous results using the same approach (10). Previous work has shown that as threshold is decreased and cluster size is appropriately increased, the rate of false positives remains stable (49).

### 2.7 Correlation analyses

We performed whole brain correlation analyses to determine whether activities in regions of significant group difference that correlated with SSRT in TD and replication groups, also correlated with SSRT, or with inattentive, hyperactive and total symptom scores in the ADHD group. These correlations were performed on all phases of activity using the following event types and contrasts: 1) warning phases (Fixate); 2) response phases ((X+O)/2); 3) reactive inhibition (SI-(X+O)/2); 4) error detection (Detect); 5) post-error slowing (PES). Statistics for slope (B1) term estimates from correlation analyses were distributed as a t* statistic with 12 degrees of freedom. All correlation maps were whole brain corrected as described above, and inspected for significant clusters in regions that exhibited significant group differences. In TD, only SSRT correlations that replicated in healthy young adults were reported.

## Results

### Performance

The 14 TD (mean (±SD) age 15.4 ±1.6) and 14 ADHD (age 13.7 ±2.1) adolescents showed significant age difference (1.71 years, p = 0.024). TD and ADHD groups showed no significant difference in post-error slowing (TD 23.9 ±35.5 ms; ADHD 10.9 ±31.2 ms; p = 0.31), go reaction time (TD 566 ±116 ms; ADHD 663.2 ±155 ms; p=0.072), percent correct go-responses (TD 97.86 ± 2.98%, ADHD 96.79 ± 4.74%, p=0.47) or the percent of successful stop trials (TD 51.41 ± 2.68%, ADHD 52.41 ± 3.66%, p=0.41). The only behavioral difference was in stop signal reaction time, which was longer (35.5 ms, p = 0.039) in the ADHD (233 ±51.0 ms) than in the TD group (198 ±33.0 ms). Behavioral performance in the replication sample was in normal range (RT = 538.5 ± 92.0 ms; SSRT = 218.3 ± 38.2 ms; post-error slowing = 12.8 ± 38.6 ms; percent successful stop trials = 50.18 ± 1.34%; percent correct go reponse = 98.47 ± 1.57%).

### Group differences in activity and intersubject correlations

We aimed to determine whether regions that activate differently in TD and ADHD, and correlate with SSRT in TD, are also correlated with SSRT or are instead correlated with symptoms in ADHD. A complete list of significant group differences during each phase of activity can be found in Additional file 1, indicating whether these regions correlated with SSRT in ADHD, TD and replication groups, and with inattentive, hyperactive, or total symptom scores in ADHD. Figures of all significant correlations during each phase of activity that are not included in the main manuscript can be found in Additional file 2.

Only three regions of significant group difference were correlated with SSRT in both TD and replication groups. Activity in each of these regions instead correlated with symptoms in ADHD, and one was also oppositely correlated with SSRT (see Figure 2).

**Figure 1.**
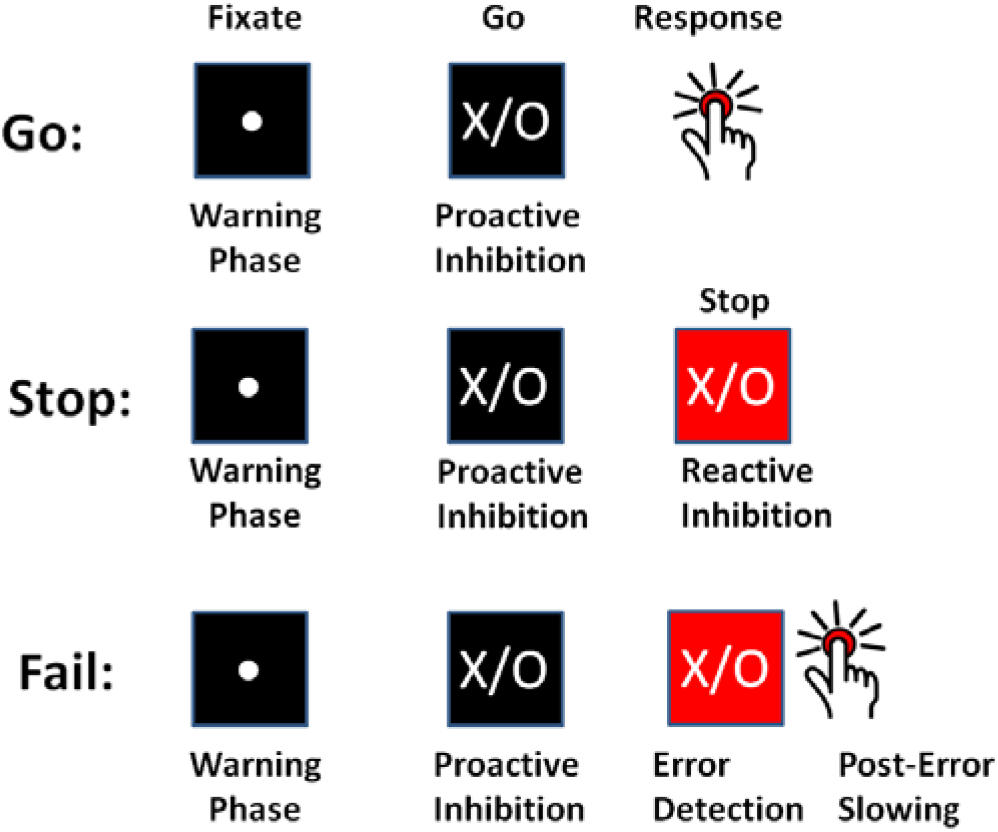
Processing stages on Go, Stop and Fail trials in the SST, all of which can contribute to inhibitory control as estimated by SSRT.

**Figure 2.**
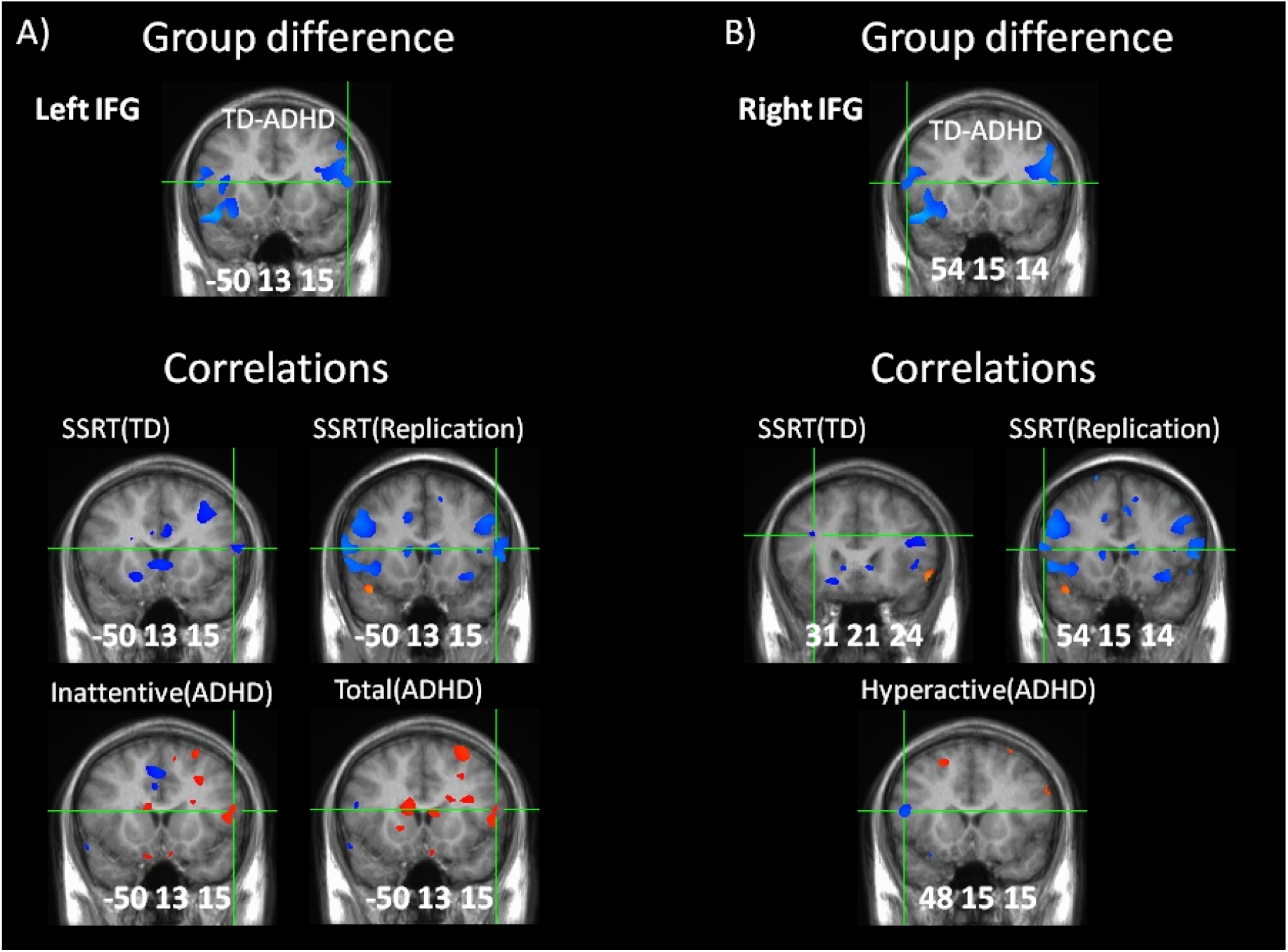
Altered inferior frontal function during error detection. A) Group difference in activity in left inferior frontal gyrus (LIFG) (top). Greater activity correlated with improved SSRT in TD and replication groups (middle) but with greater inattentive and total symptom scores in ADHD (bottom). B) Group difference in activity in right inferior frontal gyrus (RIFG) (top). Greater activity correlated with improved SSRT in TD and replication groups but with greater hyperactive symptoms in ADHD. Activation and correlation maps are whole brain corrected, and colors indicate intensity of activation (% BOLD) and correlation (r-values), red = positive activation/correlation, blue = negative activation/correlation. Locations are given in Talairach coordinates, portrayed in radiological space (left = right).

Firstly, significantly different activity (group difference z = −2.69) was present in left inferior frontal gyrus (Talairach coordinates −50, 13, 15) during error detection, the result of activation in ADHD (z = 2.76) but not in TD (z = −0.57) (see Figure 2A). Left inferior frontal activity on error detection was negatively correlated with SSRT in TD (z = −2.10) and replication groups (z = −2.08), but positively correlated with inattentive (z = 2.09) and total (z = 2.03) symptom scores in ADHD. Therefore, greater error detection activity in left inferior frontal gyrus predicted improved SSRT in TD and replication groups, whereas greater activity in ADHD instead predicted increased inattentive and total symptom scores.

Secondly, similar to left inferior frontal gyrus, significantly different activity (z = −2.69) was present in right inferior frontal gyrus (Talairach coordinates 54, 15, 14) during error detection, the result of activation in ADHD (z = 3.67) but not in TD (z = 0.81) (see Figure 2B). Right inferior frontal activity on error detection was negatively correlated with SSRT in TD (z = −2.30, at 31, 21, 24) and replication groups (z = −2.06, at 54, 15, 14) similar to right inferior frontal. However, right inferior frontal activity negatively correlated with hyperactive (z = −3.47, at 48, 15, 15) symptom scores in ADHD, in contrast with correlation of right inferior frontal with inattentive scores. Therefore, greater error detection activity in right inferior frontal gyrus predicted improved (*i.e.* shorter) SSRT in TD and replication groups, whereas greater activity in ADHD instead predicted lower hyperactive scores.

Thirdly, significantly different activity (z = −2.29) was present in the hypothalamus (Talairach coordinates −4, −5, −6) during post-error slowing, the result of deactivation in TD (z = −2.82) but not in ADHD (z = 0.82) (see Figure 3). Hypothalamus activity on post-error slowing was correlated with SSRT in TD (z = 2.50, at 4, −6, −9) and replication groups (z = 2.33, at 3, −3, −10), but correlated with inattentive (z = 2.67, at 0, −1, −11) symptom scores, and negatively correlated with SSRT (z = −2.08, at 1, −3, −5) in ADHD. Therefore, greater deactivation in the hypothalamus during post-error slowing predicted improved (*i.e.* shorter) SSRT in TD and replication groups, but instead predicted lower inattentive scores and worse (*i.e.* longer) SSRT in ADHD.

**Figure 3.**
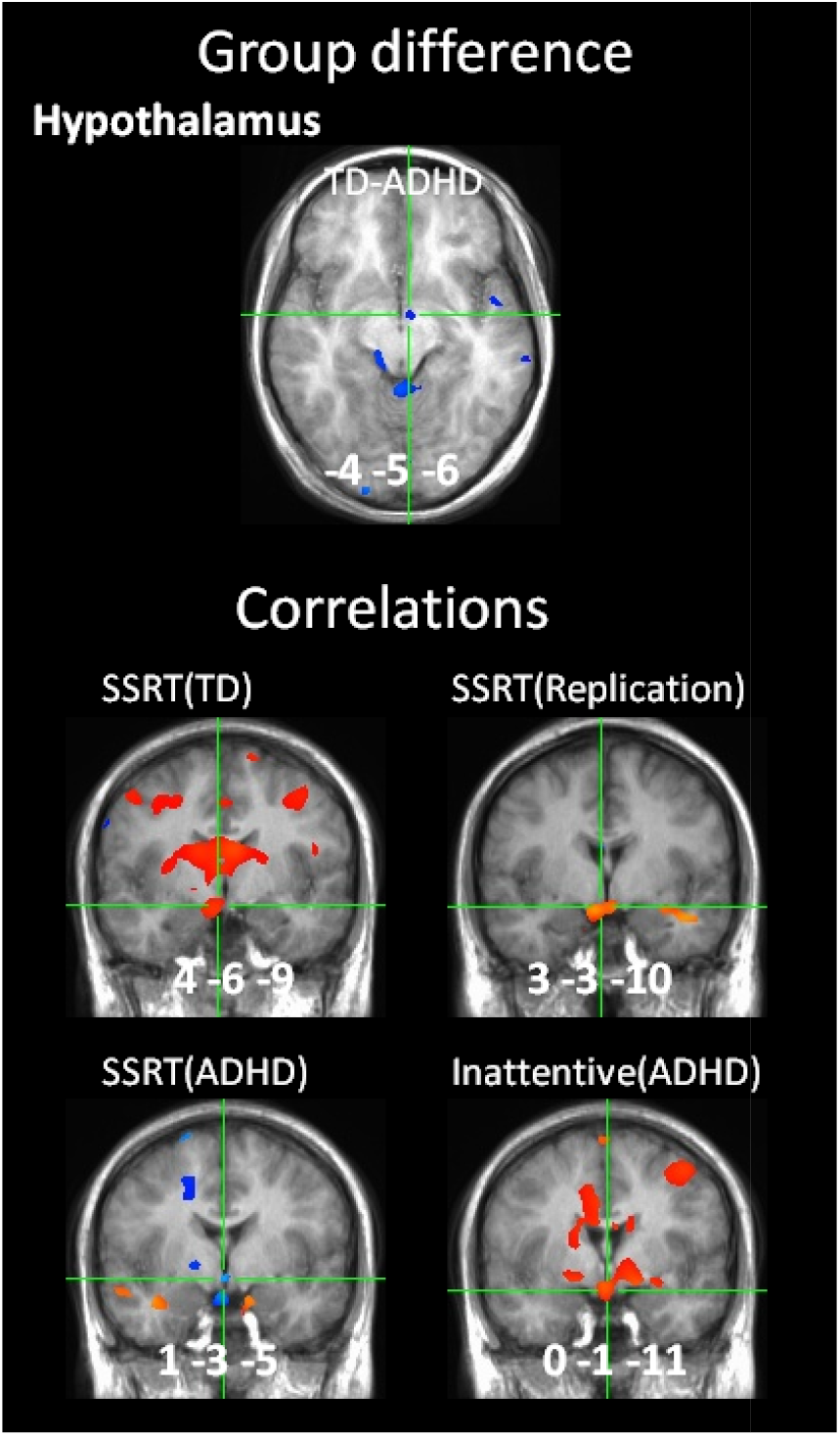
Altered hypothalamic function during post-error slowing. Group difference in activity during post-error slowing (top). Greater deactivation of the hypothalamus correlated with improved SSRT in TD and replication groups (middle), but with worse SSRT and greater inattentive symptoms in ADHD (bottom). Activation and correlation maps are whole brain corrected, and colors indicate intensity of activation (% BOLD) and correlation (r-values), red = positive activation/correlation, blue = negative activation/ correlation. Locations are given in Talairach coordinates, portrayed in radiological space (left =right).

In this study, we are primarily interested in regions of significant group difference that correlate with SSRT in both TD and replication groups, as replication provides a high level of confidence that these activities truly reflect task-directed processing in healthy subjects, and provide a meaningful baseline for comparison with ADHD. However, several brain regions that exhibited opposite activities in ADHD and TD groups, noted in the introduction, were correlated with ADHD symptoms during various phases of activity, but were not correlated with SSRT in TD and replication groups. These correlations in ADHD adolescents need to be validated by future replication, and are therefore included in Additional files 1 and 2. Some potentially important correlations in ADHD included: 1) Response phase activities in right inferior frontal gyrus, which activated in TD but deactivated in ADHD, and default mode related anterior cingulate, which deactivated in TD but activated in ADHD, both correlated with increased inattentive and hyperactive symptom scores (see Supplementary Table 2 in Additional file 1 and Supplementary Figure 2 in Additional file 2). 2) Ventral striatum, which oppositely activated in ADHD compared to TD during post-error slowing, also activated during error detection and correlated with increased SSRT and greater hyperactive symptom scores in ADHD (Supplementary table 4 in Additional file 1 and Figure 4 in Additional file 2); 3) Amygdala activity, which was heightened in ADHD during errors, correlated with inattentive symptoms during warning (see Supplementary Table 1 in Additional file 1 and Supplementary Figure 6 in Additional file 2), response (see Supplementary Table 2 in Additional file 1 and Supplementary Figure 2 in Additional file 2), error detection (see Supplementary Table 4 in Additional file 1 and Supplementary Figure 4 in Additional file 2) and post-error slowing (Supplementary Table 5 in Additional file 1 and Supplementary Figure 5 in Additional file 2) phases. 4) Cholinergic basal forebrain activity correlated with inattentive symptoms during warning (Supplementary Table 1 in Additional file 1 and Supplementary Figure 1in Additional file 2), response (Supplementary Table 2 in Additional file 1 and Supplementary Figure 2 in Additional file 2), reactive inhibition (Supplementary Table 3 in Additional file 1 and Supplementary Figure 3 in Additional file 2) and post-error slowing phases (Supplementary Table 5 in Additional file 1 and Supplementary Figure 5 in Additional file s), whereas heightened noradrenergic locus coeruleus activity correlated with hyperactive symptom scores during response (Supplementary Table 2 in Additional file 1 and Supplementary Figure 2 in Additional file 2) and post-error slowing (Supplementary Table 5 in Additional file 1 and Supplementary Figure 5 in Additional file 2) phases. 5) Hypothalamus activity, which was elevated in ADHD during post-error slowing, correlated with inattentive symptoms during warning (Supplementary Table 1 in Additional file 1 and Supplementary Figure 1 in Additional file 2) and error detection (Supplementary Table 4 in Additional file 1 and Supplementary Figure 4 in Additional file 2) phases.

## Discussion

We wanted to compare two hypotheses for the observed difference in BOLD between ADHD and TD. The conventional interpretation is that under- and over-activations in ADHD reflect weak or compensatory versions of normal function. If so, then activities in TD that correlate with task performance indexed by SSRT should also correlate with SSRT in ADHD. We proposed an alternative interpretation, that BOLD differences might instead reflect the same neural processing resources being directed toward non task-related goals, such as managing inattentive and hyperactive-impulsive symptoms associated with ADHD. This hypothesis assumes that inattentive and hyperactive symptoms do not simply reflect degraded versions of normal cognitive control, but rather alternate forms of active attention and responding that require support from the same neural processing resources that are required for normal task-directed performance, and therefore likely interfere with normal task-directed performance. If so, then activities that correlate with task performance in TD should instead correlate with inattentive and/or hyperactive symptom scores in ADHD. This alternative hypothesis has not previously been tested or explored. We tested these competing hypotheses by performing correlation analyses on all phases of activity in the SST in TD and ADHD adolescents performing the SST. Results were consistent with our alternate hypothesis that altered activities in regions that correlate with SSRT in TD instead correlate with symptoms in ADHD. The ability to test this novel hypothesis can provide a better understanding of how ADHD symptoms arise from altered neural function and subjective experience of situations that require cognitive control, such as in the SST.

The current observation of correlations with SSRT in TD but with symptoms in ADHD during error detection and post-error slowing, but not during other phases, emphasizes the need for separating and investigating these phases of activity in the SST. Many neuroimaging studies of the SST have focused on reactive inhibition activity on successful stop trials, often subtracting activity on failed stop trials based on the rationale that this contrast controls for the appearance of stop signals when it in fact confounds reactive inhibition with error processing activity. Our results are consistent with ADHD subjects exhibiting categorical differences on failed stop trials compared to TD. Understanding these categorical differences is essential for an improved understanding of ADHD and for the development of effective therapeutic interventions.

The current results might help to explain why neuroimaging studies of ADHD continue to have low effect sizes (~30-50% overlap (50,51)) despite persistent attempts to enhance these effect sizes by increasing sample size, splitting subjects by diagnostic subtypes or other individual differences, and isolating finer subcomponents of normal cognitive control. Our findings demonstrate that the important neural differences in ADHD are not simply quantitative differences in the magnitude of regional activities as much as in the nature of integrated function, and the goals toward which integrated function is directed. Overall, our results are consistent with altered activity in ADHD reflecting processing resources that normally support task-directed behavior instead being used to support or suppress wandering attention and hyperactive-impulsive behaviors associated with ADHD.

### Altered activities in regions that predict SSRT in TD, instead predict symptoms in ADHD

We found that only three significant group differences in activity were correlated with SSRT in TD and a replication sample of healthy adults: bilateral inferior frontal gyri during error detection and hypothalamus during post-error slowing. Consistent with our alternate hypothesis, all of these activities were instead correlated with symptoms in ADHD, and one of these (hypothalamus) was also oppositely correlated with SSRT compared to TD and replication groups.

It is noteworthy that activities that predict SSRT in TD but symptoms in ADHD were apparent during errors and not other phases of the task. Errors are important moments for integrating information that becomes available upon outcomes, and our results point to this level of integration being fundamentally altered in ADHD. Further, identifying these effects requires the separation of error detection from post-error slowing, which has not been done in any previous study. Our results point to the need for separating error detection from post-error slowing in the SST, because the functional roles of these two stages of processing, although tightly related, are highly distinct; error detection must interrupt ongoing processing and initiate the operating conditions necessary for adjustment, whereas post-error slowing must actually carry out the adjustment. Further, we showed in previous work (10,46) that activities in reinforcement learning related structures exhibited opposite activity (*e.g.* deactivation then activation) during error detection compared to post-error slowing, which would be lost to approaches that do not separate these phases of activity.

### Inferior frontal correlations during error detection

Negative correlation of left and right inferior frontal activity during error detection with SSRT in TD and replication groups suggests that healthy subjects who activate inferior frontal gyri more strongly at moments when errors are detected have shorter SSRT. By contrast, activation of left inferior frontal gyrus involved in interference suppression (35) correlated with greater inattention in ADHD, while activation of right inferior frontal gyrus, directly involved in response inhibition (3), correlated with less hyperactivity. These results are consistent with: 1) inhibitory control processing (right inferior frontal activity), which normally improves SSRT by supporting the inhibition of task-related responses that need to be cancelled, instead playing an active role in the suppression of impulsive behavior, and 2) interference suppression processing (left inferior frontal activity), which normally improves SSRT by suppressing forms of processing that interfere with task-directed processing in healthy subjects, instead playing an active role in supporting wandering attention in ADHD.

These findings clearly demonstrate that activity differences in ADHD do not simply reflect the over- or under-functioning of component processes, but the integration of these processing resources toward the management of symptoms instead of toward the task. Correlation of left inferior frontal activity with inattention is also consistent with our initial speculation, noted in the introduction, that inattentive and hyperactive-impulsive states associated with ADHD might cause task-related information to be targeted and processed in the brain as noise rather than signal. Most current models would hold that ADHD involves degraded neural processing of agreed upon signals and noises, rather than a different mapping of events into signals and noises. However, if task-related stimuli are not being processed by the brain as signals but rather as noise, then interventions that can strengthen signal to noise in the context of normal function might have little or potentially opposite effects on information that is being processed as noise. Although stimulant medications increase signal to noise of task-directed attention and performance in both healthy subjects and those with ADHD, our findings indicate that improvements in ADHD might be more due to a shift in the susceptibility of attentional networks to automatically process task-directed stimuli as signals rather than noises. The current approach could be used to test whether stimulant medications cause modulations in the intensity of altered activities in ADHD, or a shift toward relatively normal processing.

The importance of inferior frontal activities at moments that errors are detected might signify the central role of the efference copy of ongoing events in striatal learning. The term efference copy refers to the convergence of signals representing context, action and reward in the striatum that are necessary for reinforcement learning (52–56) In combination with our internal models, efference copies enable the brain to predict the effects of an action. Efference copies are integral to disambiguating self- from non-self generated sensation, which is central to the perception of willful behavior, and alterations in efference copy processing are thought to underlie the sense of alien control that can occur in schizophrenia (57,58). Alterations in efference copy in ADHD could drive inattentive and hyperactive-impulsive symptoms in ADHD by affecting the ability to appropriately predict and perceive the effects of actions.

Our results indicate that the efference copy of events at moments that errors are detected, which reflect the goals toward which neural processing is directed, exert a dominant influence on overall task performance in healthy subjects. However, in ADHD, altered inferior frontal activity does not likely reflect the degree of task-directed efference copy or reinforcement learning. Rather, inferior frontal activities that normally contribute to the efference copy of task-directed processing are instead appear to be engaged in managing inattentive and hyperactive symptoms, and thus cannot provide any task-related efference copy. If we cannot evoke an image (*i.e.* neural representation) of what happened leading up to the error, then there is no leverage for effective adjustment in the brief time window allowed for reinforcement learning. Therefore, therapeutic interventions aimed at strengthening such non-occurring processes would be expected to have little impact.

### Hypothalamus correlations during post-error slowing

Post-error slowing activity in the hypothalamus correlated with inattentive symptoms in ADHD, and was the only replicated correlation with SSRT in TD and healthy young adults that also oppositely correlated with SSRT in ADHD. The hypothalamus is a cognitive-motivational interface (30,31), which orchestrates the integration of distributed neural processing toward unified goals. Correlation of post-error slowing activity in the hypothalamus with inattentive symptoms in ADHD, and opposite correlation with SSRT compared to TD, is consistent with the hypothalamus orchestrating an integration of distributed neural processing in ADHD that supports wandering attention while simultaneously suppressing task-directed processing that would interfere with wandering attention.

The hypothalamus plays a central role in mobilizing and unifying distributed processing in the brain toward specific goals (food-seeking, rest, response to threats, *etc*.) based on internal states (hunger, thirst, stress etc) (30–32). However, supporting biological processes associated with one goal must also be associated with suppressing processes associated with other goals that compete for overlapping but limited resources. For example, in response to elevated levels of stress hormones or to the heightened activities in the amygdala and neurotransmitter nuclei that we observed in ADHD on errors (10), the hypothalamus can orchestrate an integration of distributed neural processing toward immediate survival, while suppressing the allocation of resources toward growth and development (reviewed in (32)), such as incremental reinforcement learning.

Here we found heightened amygdala activity on post-error slowing correlated with greater inattentive symptoms in ADHD. We also found that amygdala activity correlated with inattentive symptom scores in ADHD during all phases of the SST except reactive inhibition phases on successful stop trials. In a previous paper (10) we showed that heightened amygdala activity on errors likely drives limbic-motor interfacing conditions in the striatum, which would prevent the kind of ventral to dorsal thresholding function necessary for imposing reinforcement learning effects on distributed cortical networks (20).

Rather than simply reflecting a degraded form of normal reinforcement learning on errors in ADHD, altered striatal function in the form of limbic-motor interfacing is in fact harmonious with the absence of an efference copy required for reinforcement learning. In other words, our results are consistent with an entirely different integration of distributed processing in ADHD that is not amenable to the reinforcement learning models that appropriately describe neural function in TD. Despite the similarity of stimuli and outcomes (*e.g.* successful and failed stop trials) in both TD and ADHD groups when performing the SST, our results point to categorically different neural representations and therefore subjective experience of the same task-related stimuli and events in ADHD compared to TD.

In addition to the kind of altered integration of distributed neural processing that would be caused by heightened amygdala activity, heightened activity and altered correlations among neurotransmitter nuclei that compete for control of dopamine (10) would also support a different integration of available information on errors in ADHD. For example, heightened activities in locus coeruleus and medial septal nuclei that we found in ADHD during post-error slowing (10) would drive externally directed attention and learning about context instead of the internally directed attention required for reinforcement learning from feedback (23–26,59–61). Here we found that all the neurotransmitter nuclei that exhibited heightened activity in ADHD during post-error slowing (*i.e.* locus coeruleus, medial septal and raphe nuclei) also exhibited correlations with symptoms during various phases of activity. Given their potentially important role in driving ADHD symptoms, these correlations should be validated by replication in future work to help identify the most promising targets for interventions.

### Implications for neurocognitive models and therapeutic interventions

Neural processing resources that are directed toward the regulation of inattentive and hyperactive symptoms cannot simultaneously be used for task performance. Therefore, therapeutic interventions that attempt to directly enhance a component function that improves task performance in healthy subjects might only affect the expression of symptoms in ADHD without affecting task-directed behavioral control in general. It remains to be seen whether interventions that alleviate symptoms would cause a shift towards relatively normal functioning in ADHD. This is an important distinction for monitoring the effects and predicting outcomes of therapeutic interventions.

The current approach can test whether therapeutic interventions cause a shift toward relatively normal function in several ways, including: 1) testing whether activities that previously correlated with symptoms (*e.g.* right prefrontal and default mode networks during response phases, bilateral inferior frontal gyri during error detection, and hypothalamus during post-error slowing) become correlated with SSRT in ADHD as they do in TD adolescents, 2) testing whether therapy-induced reduction in amygdala activity causes a transition from limbic-motor interfacing to reinforcement learning related thresholding influences in the striatum on errors, 3) testing for a normalization of the competition for control of dopamine evident in normalized activities and correlations in neurotransmitter nuclei on errors. The use of such measures can improve therapeutic interventions and the neurocognitive models used to inform them.

### Limitations

Several methodological limitations of this study that restrict the interpretations of results and point to future work necessary for validating and extending the current work. Firstly, the hypothalamus and neurotransmitter nuclei identified here are small and in the brainstem, which is generally associated with increased susceptibility artefacts. However, we are confident that activities and correlations in these nuclei reflect true positives and not artefact-related noise, because they are all in specific locations that are relatively free of susceptibility artefact (62,63), and all of these regions exhibited replicated activities and inter-correlations of activity in TD and healthy young adults, and distinct activities and correlations in ADHD (10). Further, post-error slowing activity in the hypothalamus exhibited replicated correlation with SSRT in TD and healthy young adults in the current analysis. Secondly, given the small sample size and conservative subject exclusion criteria, combined with the low level of replication between SSRT correlations in TD and replication groups, ADHD correlations with SSRT and symptoms reported here are in need of replication in future studies to better determine their true validity.

Thirdly, the replication sample of healthy young adults used here is from the placebo condition of a methylphenidate study, and so might not capture SSRT correlations that would normally be replicable in the absence of placebo effects. Fourthly, SSRT and ADHD symptoms were correlated with event-related BOLD responses, but might better correlate with inter-regional connectivity estimates using time series analyses such as dynamic causal modelling (64). However, the current event related approach would still be necessary for the identification of seed locations for performing time series analyses.

## Conclusions

Our results support an interpretation that differs from the usual one which holds that altered BOLD responses in ADHD compared to TD arise from relatively weak or compensatory versions of normal task-related processing. Instead, our results are consistent with alerting stimuli that normally cause the preparation and adjustment of task-directed processing instead engaging processes involved in mediating inattentive and hyperactive symptoms, and partially in direct opposition to task-directed processing. The correlation analyses performed here demonstrate that activities during errors and preparatory periods like warning and response phases that precede the appearance of stop signals, and not those during reactive inhibition, are most strongly predictive of ADHD symptoms and of overall inhibitory control estimated by the SSRT, highlighting the importance of separating these phases of activity. A better structural understanding of the altered neural representations of task-related stimuli and events, which underlie altered cognitive control, can help us move beyond deficit based theories and ask how to gain some leverage on the integration of component processes that would be necessary for more effective therapeutic interventions.

## List of abbreviations

ADHD: Attention deficit hyperactivity disorder
AFNI: Analysis of functional neuroimages
ANOVA: Analysis of variance
BOLD: Blood-oxygen-level-dependent
DSM-5: Diagnostic and statistical manual of mental disorders, 5^th^ edition
FOV: Field of view
fMRI: Functional magnetic resonance imaging
GRE-EPI: Gradient recalled echo-echo planar imaging
HRF: Hemodynamic response function
IQ: Intelligence quotient
ITI: Inter-trial interval
MRI: Magnetic resonance imaging
ODD: Oppositional defiant disorder
PES: Post-error slowing
PICS-IV: Parent interview for child symptoms
RT: Reaction time
SD: Standard deviation
SI: Successful inhibition
SPGR: Spoiled gradient recalled
SSRT: Stop signal reaction time
SST: Stop signal task
TD: Typically developing
TE: Echo time
TR: Repetition time
WISC: Wechsler intelligence scale for children

## Declarations

### Ethics approval and consent to participate

Subjects gave informed, written consent and the study was approved by the Hospital for Sick Children institutional research ethics board. Written informed consent was obtained from the parents of all participants under the age of 16.

### Consent for publication

Not applicable.

### Availability of data and materials

The datasets supporting the conclusions of this article are available in the Mendeley Data repository, https://data.mendeley.com/datasets/x6shbht399/draft?a=25d304c3-ded9-42e4-97c7-7f545359272a.

### Authors’ contributions

AC designed the study, conducted analyses, and wrote and edited the manuscript. RS supervised study design, provided funding, and supervised and edited the manuscript. Both authors read and approved the final manuscript.

