## Supplementary Tables for "BOLD differences normally attributed to inhibitory control predict symptoms, not task-directed inhibitory control in ADHD"

**Additional file 1 - Supplementary Tables of group BOLD differences and correlations with SSRT and ADHD symptoms**

**Warning phase (Fixate)**

Group differences in activity during warning phases are listed in Supplementary Table 1. Group differences in frontoparietal regions were the result of TD but not ADHD pre-activating task-related networks, while differences in anterior cingulate were the result of deactivation in ADHD but not TD. By contrast, differences in the insula and medial septal nuclei were the result of activation in ADHD and deactivation in TD. None of these regions exhibited significant correlations with SSRT or ADHD symptoms. The hypothalamus significantly deactivated on the left in TD and on the right in ADHD, and greater deactivation in both of these locations correlated with fewer inattentive symptoms in ADHD. Further, despite an absence of significant group differences in activity, positive correlations with inattentive symptoms were also present in basal forebrain, raphe nucleus and right amygdala in the ADHD group. Group difference in activity of left postcentral gyrus was the result of activation in TD and subthreshold deactivation in ADHD, and greater deactivation in ADHD correlated with shorter SSRT.

Supplementary Table 1. Group differences and behavior/symptom correlations during warning phase (Fixate). Structures, locations with Brodmann areas, and Z scores for TD, ADHD and group difference are portrayed. Correlations with inattentive (IN), hyperactive (HI) and total (TOT) symptom scores are in brackets next to Z scores. In TD, only correlations that were also present in our replication sample of healthy young adults are reported. L = left, R = right, hTH = hypothalamus, Cb = cerebellum, rACC = rostral anterior cingulate cortex, ACC = anterior cingulate cortex, MTG = middle temporal gyrus, SFG = superior frontal gyrus, IFG = inferior frontal gyrus, LIP = lateral inferior parietal, MFG = middle frontal gyrus.

| Structure | Location | TD Z | ADHD Z | Diff Z |
| --- | --- | --- | --- | --- |
| L hypothalamus | -5 -3 -11 | -1.99 | -0.73(IN) | -0.20 |
| R ypothalamus | 2 0 -10 | -0.47 | -2.54(IN) | 1.88 |
| R cerebellum (Tonsil) | 15-47 37 | -1.30 | 1.95 | -2.38 |
| Medial Septal nuclei | 3 3 8 | -1.59 | 2.45 | -2.95 |
| L Insula 13 | -45 -21 15 | -2.90 | 2.08 | -3.51 |
| R Insula 13 | 32 7 14 | 0.05 | 3.07 | -2.80 |
| L Transverse temporal 21 | -35 -35 13 | -2.43 | 1.46 | -2.91 |
| L Fusiform 37 | -41 -63 -17 | 2.74 | -1.24 | 2.96 |
| ACC 25 | 2 5 -9 | -0.078 | -3.59 | 2.60 |
| ACC 24 | 1 27 -21 | 1.41 | -3.06 | 3.31 |
| ACC 32 | 3 44 0 | 0.36 | -3.02 | 2.54 |
| L middle temporal 22 | -58 -13 -4 | 1.41 | -2.62 | 2.95 |
| R middle temporal 22 | 59 -25 -1 | 1.66 | -2.45 | 2.89 |
| L Insula 47 | -44 15 0 | 2.45 | -0.90 | 2.49 |
| L superior frontal 10 | -33 57 -4 | 2.40 | -0.28 | 2.19 |
| R Putamen | 25 8 3 | 2.97 | 0.34 | 2.35 |
| R inferior frontal 44 | 48 11 19 | 3.04 | -0.54 | 2.92 |
| L precentral/middle frontal | -51 1 26 | 3.71 | -1.59 | 4.02 |
| L inferior parietal 40 | -35 -43 26 | 2.01 | -2.53 | 3.25 |
| L middle frontal 6 | -23 -1 53 | 2.70 | -1.01 | 2.83 |
| L precentral 6 | -46 -2 28 | 3.22 | -1.98 | 3.90 |
| L postcentral 3 | -57 -15 30 | 2.08 | -1.81 (SSRT) | 2.75 |
| R basal forebrain | 21 14 -17 | 0.86 | 1.17 (IN, TOT) | -0.87 |
| Raphe nucleus | -1 -20 -19 | -0.83 | -0.38 (IN) | -0.55 |
| R amygdala | 29 -5 -20 | -0.41 | 1.16 (IN) | -1.01 |

**Response phase (1/2(X+O))**

Group differences in activity during response phases are listed in Supplementary Table 2. Group differences in the anterior cingulate, angular gyrus and pallidum were the result of deactivation in TD and activation in ADHD. Group differences in medial septal nuclei, locus coeruleus, parahippocampus, right inferior frontal gyrus and posterior cingulate were the result of deactivation in ADHD and sub-threshold activation in TD. In ADHD, activation of default mode related anterior cingulate and deactivation of right IFG both correlated with increased inattentive, hyperactive and total symptom scores, while deactivation of left parahippocampus and locus coeruleus correlated with greater hyperactive and total symptom scores. Symptom correlations in locus coeruleus were in different locations than peak group difference in activity. Although medial septal nuclei exhibited significant group difference, it did not correlate with ADHD symptoms or with SSRT in either group, whereas basal forebrain activity correlated with fewer inattentive and total symptom scores in ADHD, but did not exhibit significant group difference in activity, or significant activity in either group.

Supplementary Table 2. Significant group differences during response phase (½(X+O)). Structures, locations with Brodmann areas, and Z scores for TD, ADHD and group difference are portrayed. Correlations with inattentive (IN), hyperactive (HI) and total (TOT) symptom scores are in brackets next to Z scores.

| Structure | Location | TD Z | ADHD Z | Diff Z |
| --- | --- | --- | --- | --- |
| Anterior cingulate 25 | -2 7 -2 | -1.44 | 1.93 | -2.62 |
| Anterior cingulate 24 | -1 27 -1 | -1.64 | 2.44 | -3.00 |
| Anterior cingulate 32 | 1 44 -3 | -2.50 | 1.57(HI, IN, TOT) | -3.12 |
| R pallidum | 11 -4 0 | -1.52 | 1.54 | -2.33 |
| L angular gyrus 39 | -56 -59 27 | -2.00 | 3.02 | -3.72 |
| R cerebellum (tonsil) | 19 -47 -34 | -0.30 | -2.17 | 2.12 |
| L locus coeruleus | -3 -35 -16 | 0.18 | -2.26 (-HI, -TOT) | 2.12 |
| L parahippocampus 36 | -21 -31 -14 | 1.05 | -2.58(-HI, -TOT) | 3.05 |
| Medial septal nuclei | 2 3 8 | 1.03 | -2.37 | 2.70 |
| R basal forebrain | 17 12 -16 | 1.39 | -1.18 (IN, TOT) | 1.67 |
| R insula 41 | 43 -23 15 | 1.06 | -3.02 | 3.31 |
| R posterior cingulate 23 | 9 -31 23 | 1.72 | -2.24 | 3.11 |
| L posterior cingulate 23 | -7 -25 22 | 1.12 | -2.18 | 2.68 |
| R inferior frontal 46 | 36 31 12 | 0.70 | -1.90(-HI, -IN, -TOT) | 2.25 |
| R amygdala/uncus | 21 -8 -20 | 1.15 | -1.64 | 2.20 |

**Reactive inhibition phase (Stop-1/2(X+O))**

Group differences in activity during successful stop phases are listed in Supplementary Table 3. Group differences in the insula were the result of deactivation in TD and not ADHD, although greater activity in ADHD predicted greater total symptom scores. LC and bilateral SN activities were correlated with total symptom scores in ADHD. Right SN activity also correlated with greater hyperactive symptoms in ADHD. Despite an absence of significant group differences in activity, right basal forebrain correlated with greater inattentive symptoms in ADHD.

Supplementary Table 3. Significant group differences during reactive inhibition (SI-½(X+O)). Structures, locations with Brodmann areas, and Z scores for TD, ADHD and group difference are portrayed. Correlations with SSRT, inattentive (IN), hyperactive (HI) and total (TOT) symptom scores are in brackets next to Z scores. In TD, only SSRT correlations that were also present in our replication sample of healthy young adults are reported. SN = substantia nigra, RN = red nucleus, Th = thalamus, STN = subthalamic nucleus, VS = ventral striatum.

| Structure | Location | TD Z | ADHD Z | Diff Z |
| --- | --- | --- | --- | --- |
| L Claustrum | -37 -11 -3 | -1.25 | 2.13 | -2.50 |
| R Insula 13 | 44 -10 20 | -2.64 | 0.81(TOT) | -2.65 |
| L cerebellum (pyramis/tonsil) | -17 -66 -29 | 2.74 | -0.82 | 2.88 |
| Culmen | 1 -49 -12 | 2.55 | -1.88 | 3.22 |
| R locus coeruleus | 5 -36 -16 | 1.18 | 1.07 (TOT) | 0.10 |
| L SN/RN | -6 -20 -10 | 1.02 | 0.43 (TOT) | 0.49 |
| R SN/Th/RN | 9 -22 -7 | 1.18 | 1.53 (TOT) | 0.70 |
| R Basal forebrain | 17 9 -11 | 1.84 | 0.99 (IN) | 1.24 |
| L Amygdala | -21 -12 -10 | 0.46 | -0.45 (IN) | 0.64 |
| R Amygdala | 17 -5 -11 | -0.25 | 1.14 (IN) | -1.10 |
| L Amygdala | -23 -2 -12 | 0.24 | 1.16 (HI) | -0.48 |
| L middle temporal 21 | -45 -10 -13 | 3.60 | -2.35 | 3.91 |
| R SN/RN/Th | 6 -15 -8 | 2.65 | -0.89 | 2.19 |
| R SN/Th | 12 -23 -6 | 0.58 | 2.08 (HI) | -0.91 |

**Error detection phase**

Group differences in activity during error detection phases are listed in Supplementary Table 4. Left putamen was correlated with total symptom scores in ADHD. Differences in bilateral inferior frontal gyrus and parahippocampus were the result of deactivation in ADHD and not TD. Left inferior frontal gyrus was negatively correlated with SSRT in TD and replication groups, and was correlated with inattentive and total symptom scores in ADHD. Parahippocampus was correlated with hyperactive and total symptom scores in ADHD. Right inferior frontal gyrus was also negatively correlated with SSRT in TD and replication groups, and was negatively correlated with hyperactive symptoms in ADHD. Despite no significant difference in group activity, ADHD activity in left amygdala correlated with inattentive and total symptom scores, in the hypothalamus correlated negatively with inattention, and in right substantia nigra correlated negatively with inattentive and total symptom scores.

Supplementary Table 4. Significant group differences during error detection. Structures, locations with Brodmann areas, and Z scores for TD, ADHD and group difference are portrayed. Correlations with SSRT, inattentive (IN), hyperactive (HI) and total (TOT) symptom scores are in brackets next to Z scores. In TD, only SSRT correlations that were also present in our replication sample of healthy young adults are reported. SN = substantia nigra, pHPC = parahippocampus, in sem lun lob = inferior semi lunar lobule.

| Structure | Location | TD Z | ADHD Z | Diff Z |
| --- | --- | --- | --- | --- |
| R hypothalamus | 2 0 -11 | 1.29 | 0.68 (-IN) | 0.16 |
| L cerebellum (tonsil) | -27 -57 -35 | -2.67 | 0.78 | -2.95 |
| R amygdala | 24 -6 -13 | -1.61 | 2.22 | -2.78 |
| L amygdala | -18 -10 -12 | 0.42 | 0.96 (IN) | -0.52 |
| R insula 13 | 36 8 16 | -1.27 | 3.28 | -3.60 |
| L insula 13 | -36 4 17 | -1.98 | 2.20 | -2.94 |
| L inferior frontal 44 | -50 13 15 | -0.57(-SSRT) | 2.76(IN, TOT) | -2.69 |
| R inferior frontal 44 | 54 15 14 | 0.81(-SSRT) | 3.67(-HI) | -2.69 |
| R middle frontal 46 | 43 20 21 | -0.65 | 3.05 | -2.76 |
| L middle frotnal 46 | -39 16 23 | -2.69 | 1.82 | -3.08 |
| R postcentral 3 | 55 -17 32 | -1.51 | 2.01 | -2.51 |
| R putamen | 31 -16 10 | -2.55 | 2.10(TOT)(VS-DS) | -3.34 |
| R dorsal striatum | 22 -2 16 | -1.75 | 1.53 | -2.34 |
| R SN / pHPC | 13 -15 -10 | -1.78 | 0.68(-IN, -TOT) | -1.75 |
| R pHPC/pallidum | 19 -14 -3 | 0.29 | 3.36(SSRT, HI, TOT) | -2.58 |
| R cerebellum (inf sem lun lob) | 23 -70 -42 | 1.86 | -1.97 | 2.67 |
| L cerebellum (inf sem lun lob) | -30 -64 -41 | 1.84 | -1.98 | 2.48 |
| R superior frontal 10 | 20 61 0 | 1.56 | -2.24 | 2.72 |
| L superior frontal 10 (polar) | -6 65 3 | 0.34 | -3.64 | 2.81 |

**Post-error slowing phase**

Group differences in activity during post-error slowing phases are listed in Supplementary Table 5. Group difference was present in left hypothalamus, and was the result of deactivation in TD but not ADHD. Hypothalamus activity was positively correlated with SSRT in TD and replication groups, but negatively correlated with SSRT and positively correlated with inattentive symptoms in ADHD. Differences in raphe nucleus and locus coeruleus were the result of deactivation in TD and subthreshold activation in ADHD, and greater activation of locus coeruleus correlated with fewer hyperactive symptoms in ADHD. Group difference in left amygdala was the result of activation in ADHD and not TD, whereas right amygdala was correlated with inattentive and total symptom scores but did not exhibit significant group difference in activity. Activity in medial septal nuclei was significantly greater in ADHD than TD but did not correlate with SSRT or symptoms, whereas basal forebrain correlated with inattentive symptoms, but did not exhibit significant group difference in activity. Group difference was present in right geniculate, which correlated with hyperactive symptoms in ADHD. Group difference activity in right medial dorsal thalamus was the result of activation in ADHD and not TD, and correlated with longer SSRT in ADHD. Group difference activity in right inferior frontal gyrus was the result of deactivation in ADHD and subthreshold activation in TD. Greater deactivation of right inferior frontal gyrus correlated with decreased inattentive, hyperactive and total symptom scores in ADHD. Although not in locations of significant group difference, right SN and amygdala correlated with total symptoms in ADHD.

Supplementary Table 5. Significant group differences during post-error slowing. Structures, locations (Talairach corrdinates) with Brodmann areas, and Z scores for TD, ADHD and group difference are portrayed. Correlations with SSRT, inattentive (IN), hyperactive (HI) and total (TOT) symptom scores are in brackets next to Z scores. In TD, only SSRT correlations that were also present in our replication sample of healthy young adults are reported. SN = substantia nigra, RN = red nucleus, STN = subthalamic nucleus, Th(va) = ventral anterior nucleus of the thalamus, Th(md) = medial dorsal nucleus of the thalamus, preSMA = presupplementary motor area.

| Structure | Location | TD Z | ADHD Z | Diff Z |
| --- | --- | --- | --- | --- |
| L hypothalamus | -4 -5 -6 | -2.82(SSRT) | 0.82(-SSRT, IN) | -2.29 |
| Raphe nucleus | -3 -21 -17 | -1.95 | 0.72 | -2.07 |
| R SN | 10 -18 -13 | 1.47 | 0.21 (TOT) | 0.65 |
| R SN/RN/STN | 7 -17 -4 | -0.15 | 0.21 (IN) | -0.24 |
| L locus coeruleus | -4 -34 -16 | -1.95 | 1.79(-HI) | -2.54 |
| L vPallidum | -17 -6 -3 | -0.83 | 2.23 | -2.34 |
| R geniculate | 16 -23 -3 | -1.72 | 1.33(HI) | -2.18 |
| Medial septal nuclei | -2 1 2 | -1.49 | 1.87 | -2.42 |
| R Basal forebrain | 16 9 -11 | 0.19 | -0.01 (IN) | 0.17 |
| L Amygdala | -20 -2 -21 | -0.82 | 2.31 | -2.46 |
| R Amygdala | 23 -12 -11 | 0.16 | 1.26 (IN, TOT) | -0.41 |
| R orbitofrontal 11 | 23 32 -12 | -1.78 | 2.32 | -2.95 |
| L parahippocampus | -26 2 -12 | -1.10 | 2.24 | -2.56 |
| L Th(va) | -8 -7 0 | -1.58 | 1.83 | -2.41 |
| R Th(md) | -6 -107 | -0.74 | 2.11(SSRT) | -2.11 |
| L Th(md) | 8 -13 8 | -0.059 | 2.81 | -2.48 |
| L superior temporal 38 | -45 9 -20 | -1.99 | 2.44 | -3.15 |
| Bilateral Lingual 18 | -2 -77 1 | -2.19 | 1.90 | -2.93 |
| L Precuneus 7 | -8 -57 42 | -1.51 | 2.76 | -3.10 |
| L postcentral 4 | -16 -35 62 | -0.48 | 2.51 | -2.44 |
| L medial frontal (preSMA) 6 | -19 2 49 | 2.29 | 1.80 | -2.94 |
| R cerebellum (tonsil) | 32 -55 -36 | 1.73 | -1.74 | 2.42 |
| R inferior frontal 45 | 48 18 9 | 1.64 | -2.60(IN, HI, TOT) | 3.14 |

Supplementary Table 6. Significant correlations during all phases of the SST. Structures with Brodmann areas, locations (Talariach coordinates), and Z scores of correlations are portrayed. Correlations with SSRT, inattentive (IN), hyperactive (HI) and total (TOT) symptom scores are in brackets next to Z scores. In TD, only SSRT correlations that were also present in our replication sample of healthy young adults are reported. L = left, R = right, hTH = hypothalamus, pHPC = parahippocampus, SN = substantia nigra, RN = red nucleus, Th = thalamus, STN = subthalamic nucleus, rep = replication group.

| Structure | Location | TD Z | ADHD Z |
| --- | --- | --- | --- |
| **a) Warning phase** |  |  |  |
| R hypothalamus | 2 0 -11 |  | 2.07 (IN) |
| Raphe nucleus | -1 -20 -19 |  | 3.44 (IN) |
| R basal forebrain | 21 14 -17  16 13 -15 |  | 3.04 (IN)  2.43(TOT) |
| R amygdala | 29 -5 -20 |  | 3.44 (IN) |
| L postcentral 3 | -57 -15 30 |  | 2.04 (SSRT) |
| **b) Response phase** |  |  |  |
| R basal forebrain | 17 12 -16 |  | -2.47 (IN)  -1.96 (TOT) |
| L locus coeruleus | -2 -35 -16 |  | -1.93 (HI)  -1.97 (TOT) |
| L pHPC 36 | -21 -31 -14 |  | -2.57 (HI)  -2.49 (TOT) |
| Anterior cingulate 32 | 1 44 -3 |  | 2.00 (HI)  2.04 (IN)  2.95 (TOT) |
| R inferior frontal 46 | 31 28 1  30 34 12  53 29 9 |  | -3.30 (HI)  -2.23 (IN)  -2.91 (TOT) |
| **c) Reactive inhibition** |  |  |  |
| R locus coeruleus | 5 -36 -16 |  | 2.07 (TOT) |
| L SN/RN | -6 -20 -10 |  | 3.23 (TOT) |
| R SN/Th/RN | 9 -22 -7 |  | 4.24 (TOT) |
| R SN/Th | 12 -23 -6 |  | 2.28 (HI) |
| R Basal forebrain | 17 9 -11 |  | 2.69 (IN) |
| L Amygdala | -21 -12 -10 |  | 3.15 (IN) |
| R Amygdala | 17 -5 -11 |  | 2.28 (IN) |
| L Amygdala | -23 -2 -12 |  | 2.45 (HI) |
| R Insula 13 | 44 -10 20 |  | 2.37 (TOT) |
| **d) Error detection** |  |  |  |
| R hypothalamus | 2 0 -11 |  | -2.81 (IN) |
| R SN / pHPC | 13 -15 -11  13 -15 -13 |  | -2.38 (IN)  -2.02 (TOT) |
| R pHPC/pallidum | 20 -14 -3  19 -14 -3  19 -14 -3 |  | 2.17 (SSRT)  2.10 (HI)  2.04 (TOT) |
| L Amygdala | -18 -10 -12 |  | 2.48 (IN) |
| L inferior frontal 44 | -50 13 15 | -2.10 (SSRT TD)  -2.08 (SSRT rep) | 2.09 (IN)  2.03 (TOT) |
| R inferior frontal 44 | 48 15 15  31 21 24  54 15 14 | -2.30 (SSRT TD)  -2.06 (SSRT rep) | -3.47 (-HI) |
| R Putamen | 31 -16 10 |  | 2.35 (TOT) |
| **e) Post-error slowing** |  |  |  |
| hypothalamus | 4 -6 -9  3 -3 -10  1 -3 -5  0 -1 -11 | 2.50 (SSRT TD)  2.33 (SSRT rep) | -2.08 (-SSRT)  2.67 (IN) |
| R SN/pHPC | 10 -18 -13 |  | 3.56 (TOT) |
| R SN/RN/STN | 7 -17 -4 |  | 3.87 (IN) |
| L locus coeruleus | 2 -34 -18 |  | -2.06 (HI) |
| R Basal forebrain | 16 9 -11 |  | 3.37 (IN) |
| R geniculate | 16 -23 -3 |  | 2.04 (HI) |
| R amygdala | 23 -12 -11 |  | 2.66 (IN)  2.37 (TOT) |
| R Th(md) | -6 -10 7 |  | 2.23 (SSRT) |
| R inferior frontal 45 | 54 19 21  47 16 15  55 17 21 |  | 3.10 (IN)  2.98 (HI)  2.03 (TOT) |
