## Supplementary Figures for "BOLD differences normally attributed to inhibitory control predict symptoms, not task-directed inhibitory control in ADHD"

**Additional file 2 – Supplementary figures of group BOLD differences and correlations with SSRT and ADHD symptoms**

**Supplementary Figure 1.** Whole brain significant activations and correlations during warning phases. A) Hypothalamus deactivated in ADHD (left) and correlated with inattentive scores in ADHD (right). B) Cholinergic medial septal nuclei were significantly less active in ADHD compared to TD while cholinergic basal forebrain correlated with inattentive and total symptom scores in ADHD. C) Correlations with inattentive scores in raphe nucleus and right amygdala, not associated with significant group difference in activity. Locations in Talairach coordinates, portrayed in radiological space (left = right).

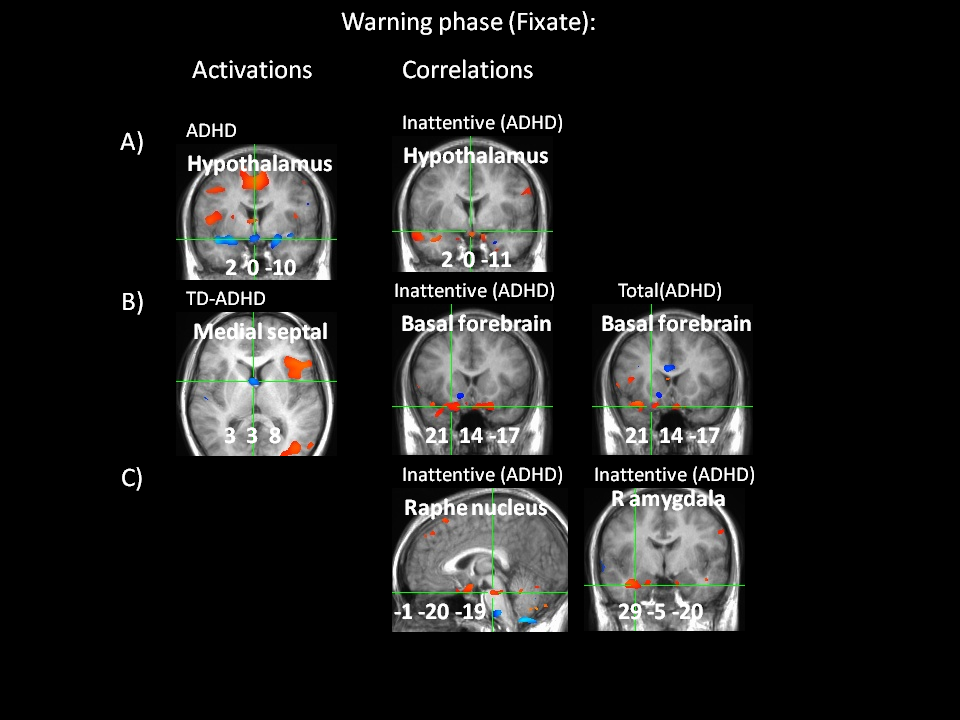

**Supplementary Figure 2.** Whole brain significant activations and correlations during response phases. A) Anterior cingulate cortex (ACC) was more active in ADHD compared to TD (left), and correlated with inattentive, hyperactive and total symptom scores in ADHD (right). B) Right inferior frontal gyrus (IFG) was less active in ADHD, and correlated with inattentive, hyperactive and total symptom scores. C) Locus coeruleus (LC) was less active in ADHD and correlated with hyperactive and total symptom scores. D) Left parahippocampus (pHPC) was less active in ADHD and correlated with hyperactive and total symptom scores. E) Medial septal nuclei were less active in ADHD and basal forebrain correlated with inattentive and total symptom scores.

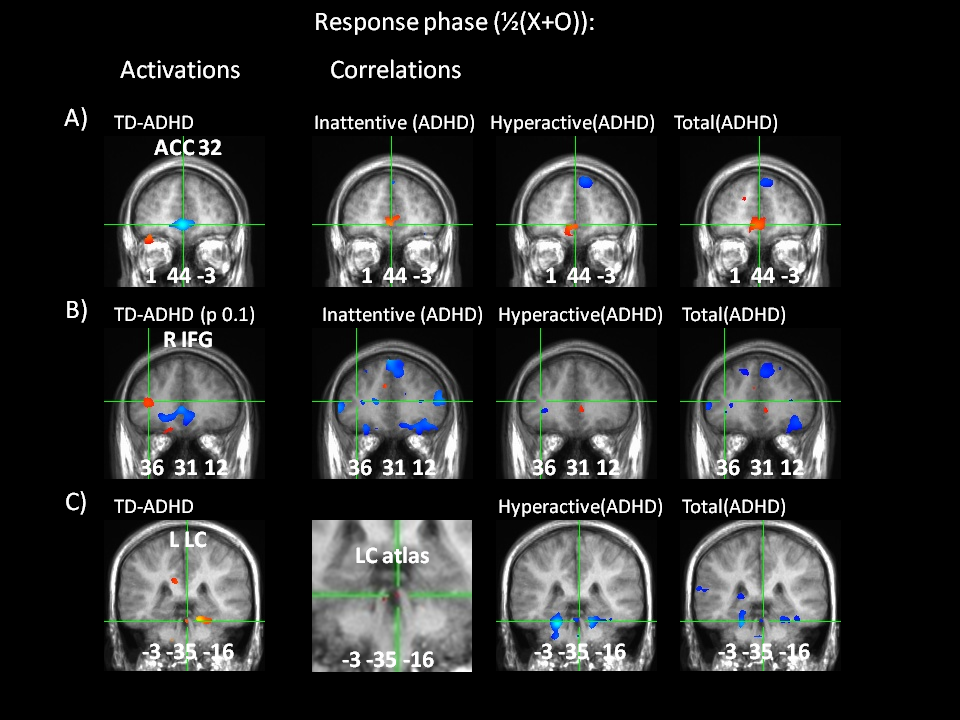

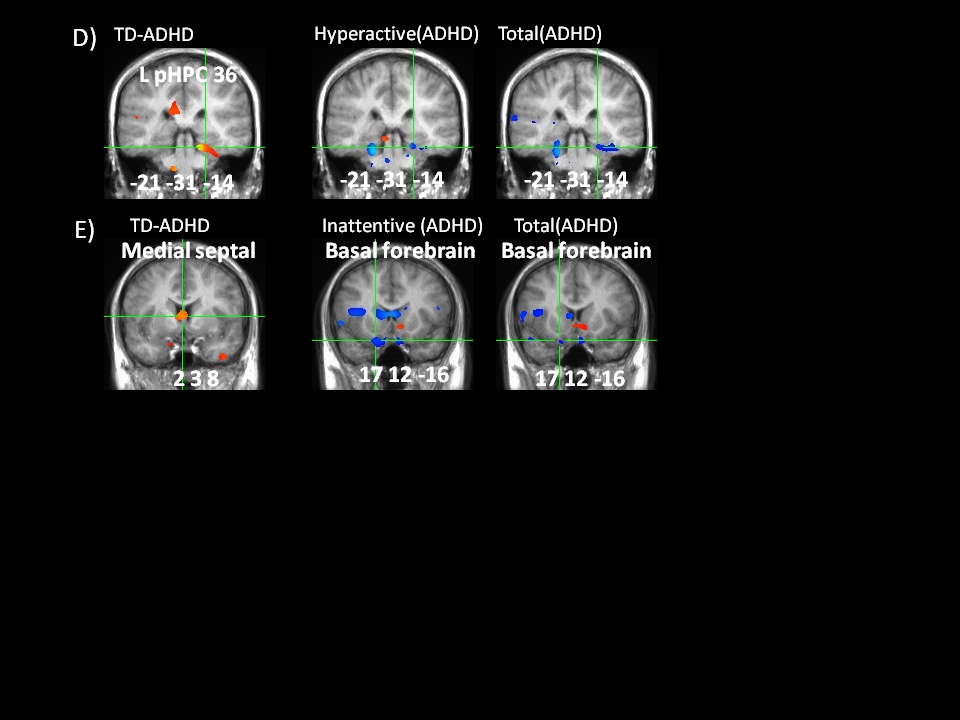

**Supplementary Figure 3.** Whole brain significant activations and correlations during reactive inhibition. A) Right insula was significantly less active in ADHD compared to TD (left), and correlated with total symptom scores in ADHD (right). B) Dorsal striatum was significantly less active in ADHD (corrected at voxelwise p=0.1) and correlated with inattentive scores. C) Other correlations not associated with significant activity or group difference in activity. LC = locus coeruleus, SN = substantia nigra, RN = red nucleus, Th = thalamus.

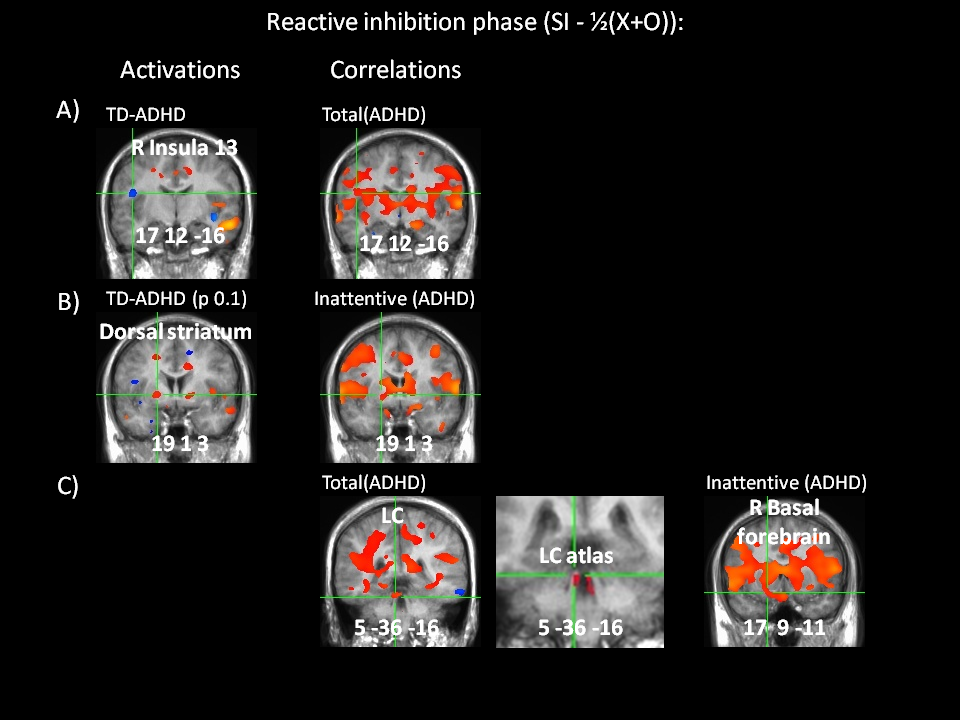

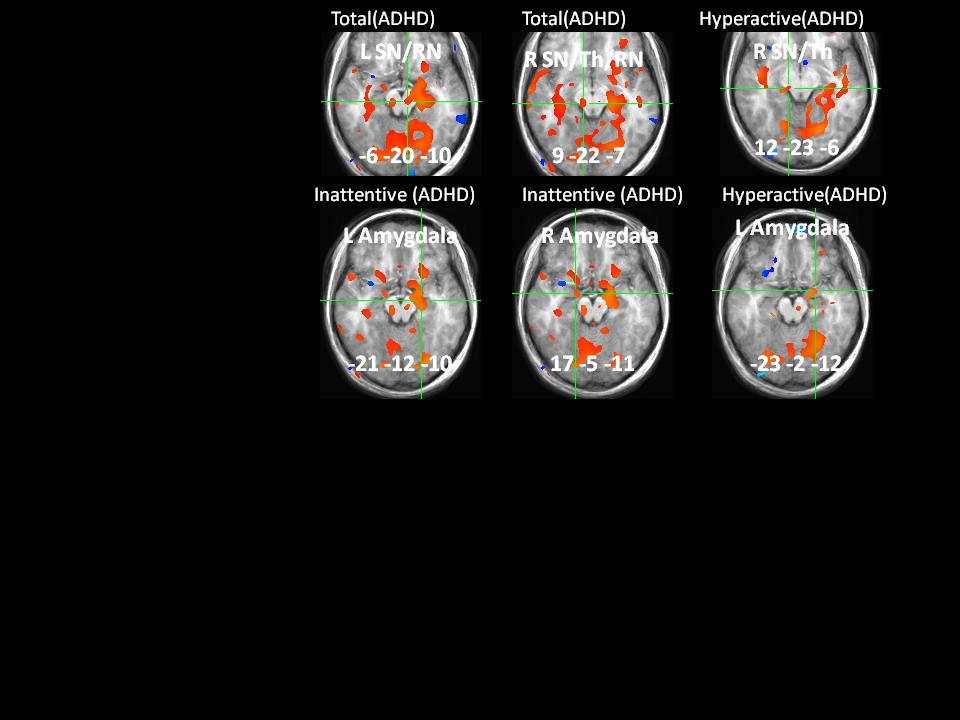

**Supplementary Figure 4.** Whole brain significant activations and correlations during error detection. A) Hypothalamus was less active in ADHD (voxelwise p=0.1) and correlated with total symptoms. B) Substantia nigra/parahippocampus (SN/pHPC) was significantly more active in ADHD and correlated with inattentive and total symptoms. C) pHPC/pallidum was more active in ADHD (voxelwise p=0.1) and correlated with SSRT, hyperactive and total symptoms in ADHD. D) Right putamen was more active in ADHD and correlated with total symptoms in ADHD. E) Right amygdala was more active in ADHD and correlated with inattentive and total symptoms.

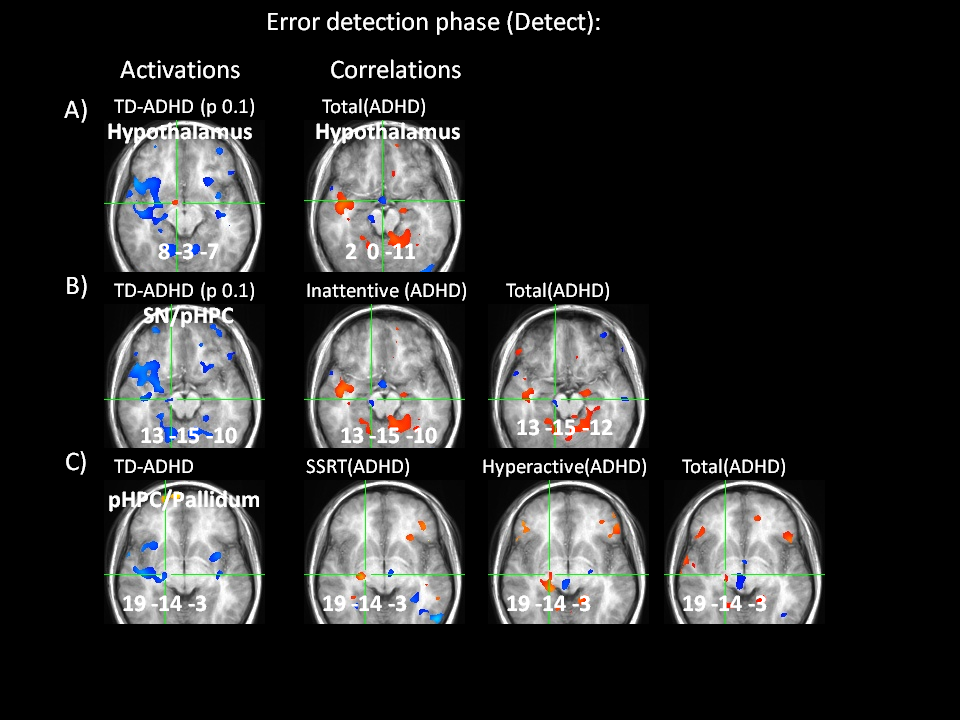

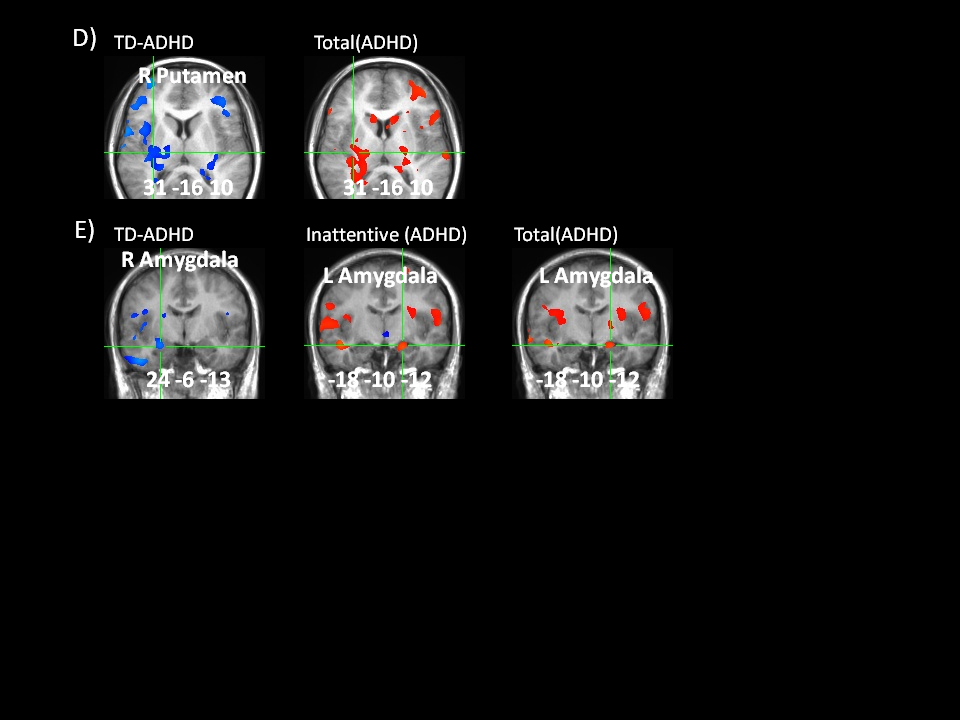

**Supplementary Figure 5.** Whole brain significant activations and correlations during post-error slowing. A) Locus coeruleus (LC) was significantly more active in ADHD (left) and correlated with hyperactive symptoms. B) Cholinergic medial septal nuclei were significantly more active in ADHD and correlated with inattentive symptoms. C) Right geniculate was more active in ADHD and correlated with hyperactive symptoms. D) Left amygdala was more active in ADHD and right amygdala correlated with total symptom scores. E) Medial dorsal thalamus (Th(MD)) was more active in ADHD and correlated with SSRT in ADHD. F) Right inferior frontal gyrus (IFG) was less active in ADHD ad correlated with inattentive, hyperactive and total symptom scores. G) Correlations not associated with significant activity or group difference. SN = substantia nigra, RN = red nucleus, STN = subthalamic nucleus, pHPC = parahippocampus.

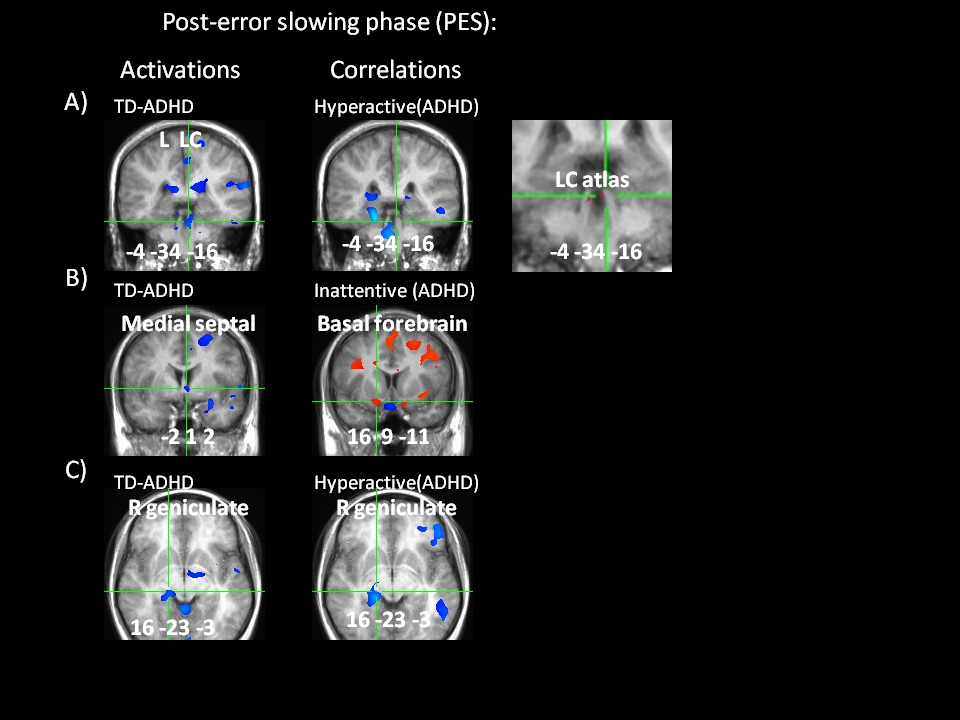

**Supplementary Figure 5 continued…**

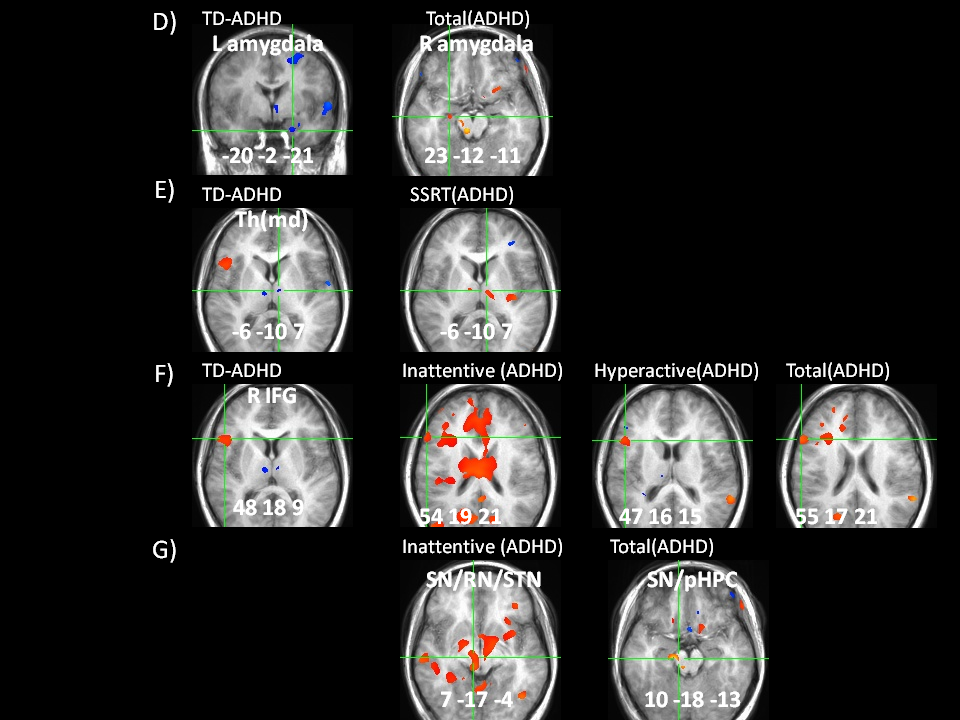
